# A Validated Method for Detection of *Bifidobacterium infantis* Bi-26^TM^ DNA in Stool by Quantitative Real-Time PCR

**DOI:** 10.1101/2025.11.26.690772

**Authors:** Markus Gödderz, Justine Sunshine, Melanie Polke, Paul Rhyne, Noel Taylor, Anthony Kiefer, Nicolas Yeung, Hilmar Wisplinghoff, Ulrich Zechner, Nicole Frahm

## Abstract

**Background:** *Bifidobacterium longum* subspecies *infantis* (*B. infantis*) is a commensal bacterium enriched in the microbiota of breastfed infants and associated with beneficial effects on infant gut health. Clinical studies evaluating *B. infantis* supplementation in infants have reported increased colonization, reduced inflammation, enhanced vaccine responsiveness, and improved growth outcomes. To support rigorous evaluation of these effects in further clinical trials, reliable and validated methods for quantifying *B. infantis* in stool are needed.

**Methods:** We validated a quantitative real-time PCR (qPCR) assay against a conserved *B. infantis* genomic target. Assay performance was evaluated using DNA from the *B. infantis* strain Bi-26^TM^, spiked infant stool pools, and matrix controls. Validation parameters included linearity, sensitivity, accuracy, within- and between-run reproducibility, recovery, and matrix interference, with acceptance criteria informed by bioanalytical best practices.

**Results:** The assay demonstrated robust performance across validation parameters. Linearity was confirmed over at least six orders of magnitude (R^2^ > 0.99; mean efficiency 96.4%), with a lower limit of quantification of 37.36 genome equivalents / reaction. Analytical sensitivity was high, with a 95% limit of detection of 4.67 genome equivalents / reaction. Accuracy across multiple stool pools met acceptance criteria, with mean deviation from nominal values of 0.39 log_10_. Reproducibility was strong, with within-run variability of 0.37% and between-run variability of 1.78%. Recovery averaged 41.3%, exceeding the predefined minimum threshold, and matrix interference was minimal.

**Conclusions:** This study reports a validated qPCR assay suitable for quantifying *B. infantis* Bi-26 DNA in stool samples collected in stabilized formats. Its robust analytical performance supports its use in clinical and translational research evaluating *B. infantis* colonization and its health impacts in early infant life.

## Introduction

*Bifidobacterium longum* subspecies *infantis* (*B. infantis*) is a commensal bacterial strain that is common in the microbiome of breastfed infants younger than 6 months of age, particularly in low-and middle-income countries (1). This strain has been shown to be an early and effective colonizer of the infant gut due to its unique array of glycosyl hydrolases, transporters, and intracellular enzymes that efficiently internalize and degrade human milk oligosaccharides (HMOs) found in breast milk (2–4). The acidic byproducts of this HMO degradation have been shown to lower gut pH, which helps suppress the outgrowth of potentially pathogenic bacteria (5–7). In addition, *B. infantis* colonization and byproducts have been associated with strengthening the gut barrier and modulating early life immune responses (8–14).

Given its potential benefits, there is considerable interest in the evaluation of *B. infantis* probiotic supplementation on infant health and development outcomes. Clinical studies across various *B. infantis* strains have shown that supplementation results in stable *Bifidobacterium* colonization and microbiome remodeling (6, 15–19), reduced gut inflammation (11), immune modulation (14), and improved responsiveness to infant vaccination (12). Clinical studies in preterm infants have shown that probiotic supplementation, often combination products including *B. infantis*, has been associated with a lower incidence of necrotizing enterocolitis (20, 21). Recently, there has been growing interest in evaluation of *B. infantis* supplementation on nutritional recovery and improved growth, especially in LIMCs where accessible and effective therapeutic options are highly limited (17, 22, 23). Notably, the SYNERGIE study, in which a *B. infantis* strain was provided for 28 days either alone or with a prebiotic to underweight infants 2 to 6 months of age in Bangladesh, demonstrated improvement in anthropometric measures including weight-for-age (WAZ) and weight-for-length Z score [WLZ] compared to placebo (17).

As clinical interest in *B. infantis* continues to expand, there is a corresponding need for accurate, sensitive, and validated methods to detect and quantify *B. infantis* levels in infant stool clinical specimens. Quantitative real-time PCR (qPCR) methods are frequently used to detect probiotic DNA in stool samples. Multiple qPCR assays have been developed to measure *B. infantis* levels in stool (17, 24–27). However, many published assays lack formal validation according to bioanalytical standards (28) with limited data on key parameters such as linearity, sensitivity, accuracy, reproducibility, recovery, and matrix interference effects. This gap highlights the need for thoroughly validated qPCR methods to support reliable detection and quantification of *B. infantis* in clinical trials.

Here, we report the validation of a qPCR method for quantifying *B. infantis* Bi-26^TM^ DNA in infant stool. The assay was developed to support clinical trials involving *B. infantis* strain Bi-26 and it may provide a valuable basis for detecting alternative *B. infantis* strains in clinical and research settings.

## Methods

### Bi-26 DNA Material

*B. infantis* strain Bi-26^TM^ (International Flavors and Fragrances, IFF) was used to generate reference standard. A total of 1.6 g of B. infantis Bi-26 powder (IFF) was resuspended in 7.5 ml of Tris-EDTA (TE) buffer (10 mM Tris, 0.1 mM EDTA, pH 8.0) and stored at −80 °C in 1 ml aliquots.

### Quality Control (QC) Stool Pool Generation

There are several commercially available collection devices containing buffers that are optimized to stabilize and preserve DNA in stool samples. For this validation we chose OMNIgene® GUT – OM-200 collection tubes (OM-200; DNA Genotek, Ontario, Canada) to ensure sample integrity. Healthy infant and adult stool samples were procured and collected in OM-200 tubes (BioIVT) and were screened for endogenous *B. infantis* and *B. longum*. Negative donors were then used to generate three different stool pools (P1, P2, P3), each containing material of at least three different individual stool samples, for a total volume of 6.8 ml per pool. To facilitate proper pipetting in case of a solid or too viscous consistency of the individual stool samples in OM-200 tubes, samples were incubated at 50 °C for 30 min according to the manufactureŕs instructions followed by brief centrifugation. The three stool pools were aliquoted and stored at −20 °C. For each stool pool, a high (1.6 ng/µl), medium (160 pg/µl) and low (5 pg/µl) positive control sample was generated (i.e. P1 High, P1 Med, P1 Low, etc.). Bi-26 DNA was directly spiked into the prepared ZR BashingBeadTM Lysis Tubes and filled up to a total sample volume of 200 µl by adding the respective stool. Unspiked (i.e. no Bi-26 DNA added) stool pools were used as negative controls in DNA extraction and subsequent experiments.

### DNA Extraction

DNA extraction of Bi-26 aliquots or stool pools was performed using the ZymoBIOMICS™ MagBead DNA kit (Zymo Research, USA) on the KingFisher Flex system (ThermoFisher Scientific, Massachusetts USA) with minor modifications (notably, switch to 800 µl lysis buffer and 200 µl sample, with additional MagBinding buffer wash). KingFisher plates were prepared with MagBinding buffer, wash buffers, and DNase/RNase-free water for elution according to the manufacturer’s instructions with the following deviations: Bead-beating tubes were filled with 800 µl lysis buffer and the sample plates were filled with 800 µl MagBinding buffer. For DNA extraction from stool pools (spiked and unspiked), each tube received 800 µl of lysis buffer, followed by bead beating in a Precellys Evolution homogenizer (Bertin Technologies, France) using the manufacturer’s recommended program. Elution was performed in 100 µl DNase/RNase-free water. DNA concentration and quality were assessed using Qubit 4 Fluorometer (quantification), NanoDrop One spectrophotometer (purity ratios; A260/280, A230/260), and Agilent 2100 Bioanalyzer (integrity profiling) according to manufacturer protocols. After extraction, eluted DNA was stored for subsequent PCR runs at 4 °C for short term storage (up to 1 week) or −80 °C for long term storage (> 1 week).

### qPCR Method

All instruments are listed in Supplementary Table 1 and all reagents are listed in Supplementary Table 2. The PCR setup and DNA dilutions were set up on ice. The qPCR assay targets an intergenic region near the phosphate acetyltransferase gene of Bi-26 and the primers were designed to detect *B. infantis* Bi-26 strain and distinguish from closely related *B. longum* species (forward (GTCACGATGTCTCCTTTGATATCAGCATG) and reverse (CCTTTTGCGTCTCCCCCG) primers, probe /5FAM/TCATTCATT/ZEN/GTAGTGGCGATCACCGTTACC/3IABkFQ/]. Primer sequences to the target gene are generally conserved across public *B. infantis* strains (data not shown). PCR mixtures contained 10 µM each of primers and probe, 5 µl nuclease-free water, 12.50 µl of TaqMan fast Advanced Master Mix 2x, and 5 µl of template DNA for a total volume of 25 µl (Table 2). DNA samples with concentrations exceeding 2 ng/µl were diluted 1:10 in TE buffer pH 8.0, eluates with DNA concentrations < 2 ng/µl were used undiluted in the PCR. Samples for validation were analyzed in technical triplicates. C1000 Thermal Cycler with CFX96 Optical Reaction Module was used with the following parameters: AmpliTaq activation at 95 °C for 20 s for 1 cycle, followed by 40 cycles of 95 °C for 3 s, 60 °C for 30 s, with final cooling cycle at 8 °C for 30 s. CFX Manager^TM^ Dx software in (Version 3.1, Bio-Rad) single threshold mode was used for quantification cycle (Cq) determination.

### B. infantis Reference Standard and Linearity

Using extracted Bi-26 DNA, a *B. infantis* reference standard with 10-fold dilution series was prepared in TE buffer (pH 8.0) using DNA LoBind tubes. The highest DNA concentration in the series was 2 ng/µl, with serial dilutions extending down to 20 fg/µl. The linearity of the *B. infantis* qPCR was evaluated by analyzing a logarithmic dilution series of *B. infantis* reference standard using a CFX96^TM^ Real-Time PCR Detection System (Bio-Rad, Colorado, USA). Each concentration was analyzed in triplicate measurements per run for three runs on three different days for a total of nine replicates per concentration. Slope and R² of semi-log regression of each PCR run were calculated. Efficiency was calculated by the following formula:

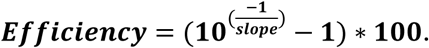

### Sensitivity

Based on the DNA sequence of *B. infantis* Bi-26 (NCBI Reference Sequence: NZ_CP054425.1) the mass of the DNA was converted to genome equivalents using the NEBioCalculator^®^ (https://nebiocalculator.neb.com/, v.1.15.4, New England Biolabs). The analytical sensitivity was determined using a dilution series of the extracted *B. infantis* Bi-26 DNA in TE buffer (10 mM Tris, 0.1 mM EDTA, pH 8.0) ranging from 37.36 genome equivalents / reaction to nominal 0.58 genome equivalents / reaction, corresponding to dilution series. Over three days, eight replicates per concentration were analyzed in qPCR for a total of 24 replicates per concentration. The results were determined by Probit analysis.

### Within-run and Between-run Reproducibility

Within-run reproducibility was assessed by analyzing High, Medium, Low, and Negative control samples from each of the three stool pools (n = 12), along with a process control containing only OMNIgene® GUT – OM-200 buffer. All extracted samples were run in triplicate. Variability was reported as the coefficient of variation (%CV). Within-run variability was calculated based on the Cq values from triplicate measurements of each sample, and total within-run variability was determined by averaging the %CV across all non-negative samples. Between-run variability was calculated using the mean Cq values from each day’s triplicate measurements, with the total between-run variability reported as the average across all non-negative samples.

### Accuracy

The accuracy of the *B. infantis* qPCR was evaluated based on the data generated during the assessment of the between-run reproducibility. Quantifications (***c_total_***/concentration) were determined by correcting the values determined by PCR (***c_PCR_***/concentration) to compensate for the PCR dilution factor ***F_Dilution_***, sample usage, lysate usage, and elution volume:

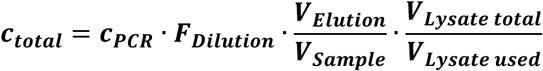

Accuracy is expressed as the logarithmic (base 10) deviation between quantified and nominal values.

The log_10_ deviation for each sample was calculated from the deviation of the mean quantified value across all six measurement days to the nominal values. Individual day values were determined from the mean value of triplicate measurements that were performed on each day. Total accuracy was calculated as the average of all (non-negative) samples.

### Recovery

The recovery rate of the assay was back-calculated based on the data generated during the assessment of between-run reproducibility. Recovery is the percentage calculated by dividing the measured value by the nominal value (the actual amount of DNA that was added to the sample prior to extraction) of the sample.

### Matrix Interference

Matrix interference was evaluated for stool samples in OMNIgene^®^ GUT – OM-200 buffer. For matrix interference experiments, ten individual stool samples collected in OMNIgene® GUT-OM-200 tubes were either spiked with a medium concentration of Bi-26 DNA (160 pg/µl) or left unspiked as negative controls. Additionally, three OM-200 buffer samples were directly spiked with Bi-26 DNA at the same concentration. Matrix interference was calculated as the log_10_ deviation between the measurement value of individual non-negative control samples to the mean value obtained from the spiked OM-200 buffer samples. The log_10_ deviation from the nominal values was calculated.

## Results

The intended use of this assay is to enable accurate quantification of *B. infantis* Bi-26 DNA levels in stool samples collected in clinical research settings, such as nutritional intervention or microbiome colonization studies. Design for this assay is based on primer set 1 from the USP monograph for *Bifidobacterium longum* subsp. *infantis* (Bi-26), enhanced by the addition of a probe for improved accuracy and amplification (29). The assay targets an intergenic region near the phosphate acetyltransferase gene of Bi-26. This assay was validated according to selected key bioanalytical performance parameters based on industry best practices, including linearity, accuracy, sensitivity, reproducibility (both within- and between-run), recovery, and matrix interference. In defining the acceptance criteria for validation success, we referred to the recommendations outlined by Hays et al. 2022 (Supplementary Table 3) (28).

We first evaluated the dilutional linearity of the assay using a *B. infantis* reference standard with DNA extracted from strain Bi-26. The linear range of the *Bifidobacterium infantis* Bi-26 DNA qPCR was shown to extend over an interval of at least six orders of magnitude ranging from 100 fg/reaction to 10 ng/reaction (Figure 1A-C). Linearity was confirmed for assay, with all regression models having R^2^ ≥ 0.99 (Figure 1D). The mean efficiency of the assay was 96.39% (Figure 1D). The lower limit of quantification (LLOQ) and upper limit of quantification (ULOQ) are defined by the lowest and highest concentrations used in this assessment, respectively. In the unit of genome equivalents / reaction the linear range of the assay is 37.36 genome equivalents/reaction (LLOQ) to 3.736 x 10^6^ genome equivalents / reaction (ULOQ). We next evaluated the sensitivity of the assay through testing an extension of the reference standard dilution series ranging from 37.36 genome equivalents / reaction to nominal 0.58 genome equivalents / reaction in 24 replicates. The analytical sensitivity of the assay is expressed as the Limit of Detection (95%) with the LOD_95_ determined to be 4.67 genome equivalents / reaction (95%-Confidence Interval: 3.39 – 8.04 genome equivalents / reaction) (Table 1).

**Figure 1:**
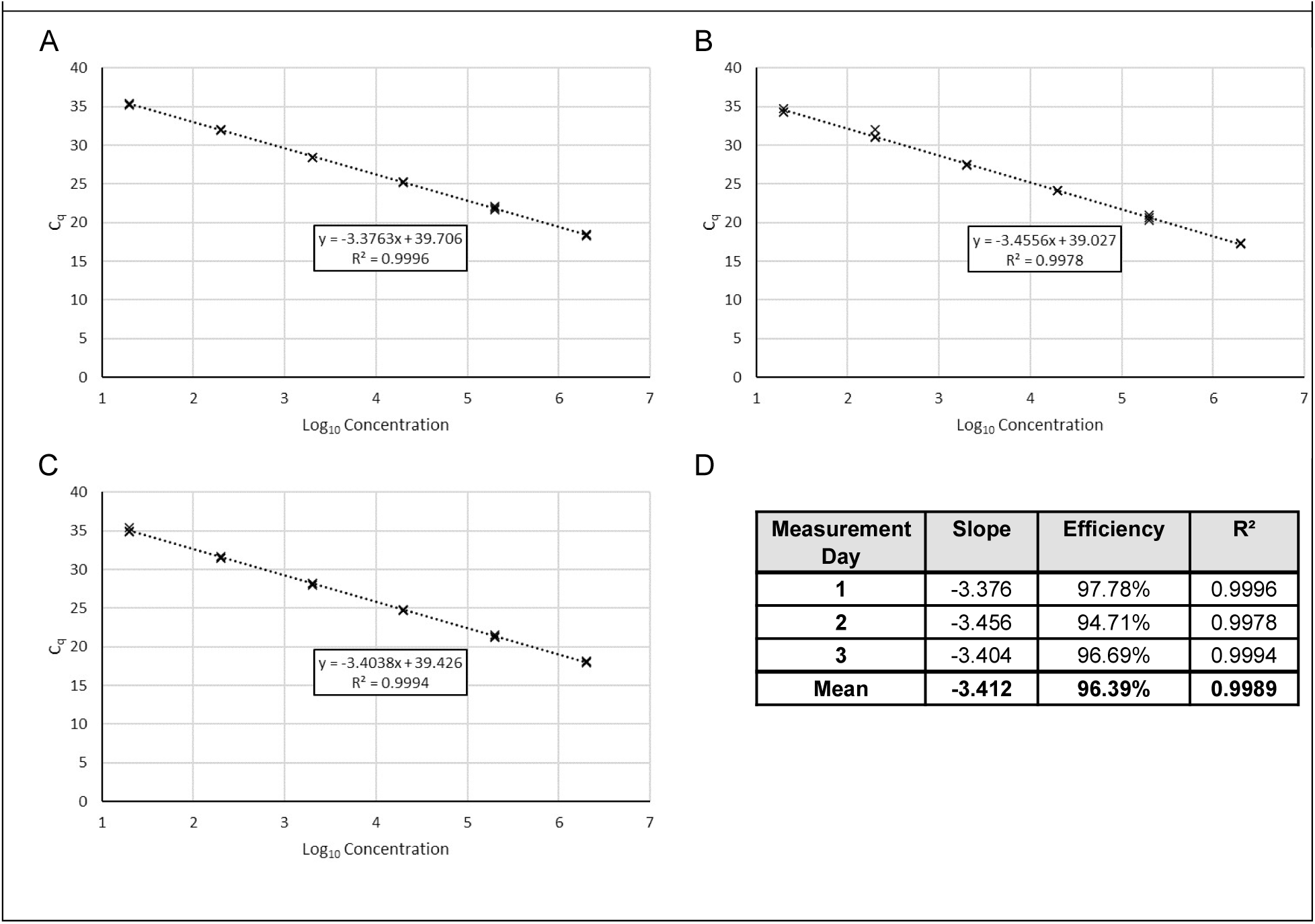
Dilutional Linearity. Semi-logarithmic display of C_q_-values vs. input concentration on A) Day 1, B) Day 2. C) Day 3. D) Summary of Slope, Efficiency, and R² of semi-log regression of each PCR run

**Table 1:** Analytical sensitivity assessment using a *B. infantis* Bi-26 DNA dilution series.

| DNA Input (fg/reaction) | DNA Input (genome equivalents/reaction) | Number of Replicates | Number of Positives | Hit Rate (%) |
| --- | --- | --- | --- | --- |
| 100 | 37.36 | 24 | 24 | 100 |
| 50 | 18.68 | 24 | 24 | 100 |
| 25 | 9.34 | 24 | 24 | 100 |
| 12.5 | 4.67 | 24 | 24 | 100 |
| 6.25 | 2.34 | 24 | 16 | 67 |
| 3.125 | 1.17 | 24 | 8 | 33 |
| 1.5625 | 0.58 | 24 | 4 | 17 |

**Table 2:** Accuracy. Accuracy reported as log_10_-deviation of mean quantification values of triplicate measurements of all control samples per day from nominal values. Sample ID includes Pool#)_(<u>H</u>igh/<u>M</u>ed/<u>L</u>ow/<u>N</u>egative/<u>OM</u> buffer only)

|  |  |  |  |  |  |  |  |  |
| --- | --- | --- | --- | --- | --- | --- | --- | --- |
| Sample ID | Spike-In Level (fg/μl) | Day 1 (fg/μl) | Day 2 (fg/μl) | Day 3 (fg/μl) | Day 4 (fg/μl) | Day 5 (fg/μl) | Day 6 (fg/μl) | Log<sub>10</sub> Deviation |
| P1_H | 1,600,000 | 305,500 | 726,150 | 773,150 | 737,900 | 620,400 | 780,200 | **0.39** |
| P2_H | 1,600,000 | 425,350 | 1,137,400 | 1,057,500 | 740,250 | 900,050 | 667,400 | **0.29** |
| P3_H | 1,600,000 | 270,250 | 848,350 | 829,550 | 615,700 | 653,300 | 789,600 | **0.38** |
| P1_M | 160,000 | 62,745 | 67,915 | 62,510 | 74,495 | 62,980 | 78,255 | **0.37** |
| P2_M | 160,000 | 68,620 | 66,035 | 98,465 | 86,715 | 62,040 | 96,115 | **0.30** |
| P3_M | 160,000 | 58,515 | 68,620 | 69,090 | 63,920 | 64,625 | 60,395 | **0.40** |
| P1_L | 5,000 | 1,332 | 1,473 | 1,739 | 1,584 | 1,986 | 1,793 | **0.48** |
| P2_L | 5,000 | 1,793 | 1,748 | 1,661 | 1,784 | 2,961 | 2,538 | **0.38** |
| P3_L | 5,000 | 1,276 | 1,901 | 1,062 | 1,664 | 1,396 | 1,725 | **0.52** |
| P1_N | 0 | ND | ND | ND | ND | ND | ND | **NA** |
| P2_N | 0 | ND | ND | ND | ND | ND | ND | **NA** |
| P3_N | 0 | ND | ND | ND | ND | ND | ND | **NA** |
| OM | 0 | ND | ND | ND | ND | ND | ND | **NA** |
|  |  |  |  |  |  |  | **Mean** | **0.39** |

Accuracy and reproducibility are critical in clinical qPCR assays to ensure true DNA quantification and consistent results within and across runs, supporting reliable comparisons across samples, subjects, and timepoints. Accuracy was calculated as described in methods and reported as the logarithmic (base 10) deviation between quantified and nominal values. Across all runs from 12 stool pools samples, the total mean trueness was determined to be 0.39 log_10_ levels, again meeting the targeted acceptance criteria of reported values being within 1 log_10_ level for ≥75% of samples (Table 2). We assessed between-run reproducibility using all three stool pools (P1, P2, P3) tested in triplicate across six days, two operators and three instruments (see methods). Within-run reproducibility was determined from the triplicate measurements of day 1 of the between-run measurements. Within-run and between-run reproducibility are reported in terms of %CV). Within-run and between-run reproducibility were acceptable with the mean within-run variability calculated to be 0.37% (Table 3) and the between-run variability determined to be 1.78% (Table 4). Both results were well-below the target acceptance criteria of <15% for within-run and <25% for between-run reproducibility.

**Table 3:** Within-run Reproducibility. Within-run variability expressed as coefficient of variation of quantification cycle values of triplicate measurements of all stool pool samples. Sample ID includes (Pool#)_(High/Med/Low/Negative)_(Day#) OM=OM buffer alone.

|  |  |  |  |  |  |
| --- | --- | --- | --- | --- | --- |
| Sample ID | Spike-In Level (fg/μl) | C<sub>q</sub> Replicate 1 | C<sub>q</sub> Replicate 2 | C<sub>q</sub> Replicate 3 | %CV |
| P1_H_D1 | 1,600,000 | 25.65 | 25.79 | 25.80 | 0.33% |
| P2_H_D1 | 1,600,000 | 25.33 | 25.23 | 25.16 | 0.34% |
| P3_H_D1 | 1,600,000 | 26.17 | 25.85 | 25.78 | 0.80% |
| P1_M_D1 | 160,000 | 28.17 | 28.18 | 28.10 | 0.15% |
| P2_M_D1 | 160,000 | 28.03 | 28.01 | 28.00 | 0.05% |
| P3_M_D1 | 160,000 | 28.30 | 28.30 | 28.17 | 0.27% |
| P1_L_D1 | 5,000 | 33.80 | 34.16 | 34.09 | 0.56% |
| P2_L_D1 | 5,000 | 33.50 | 33.47 | 33.71 | 0.39% |
| P3_L_D1 | 5,000 | 34.10 | 33.91 | 34.23 | 0.47% |
| P1_N_D1 | 0 | ND | ND | ND | NA |
| P2_N_D1 | 0 | ND | ND | ND | NA |
| P3_N_D1 | 0 | ND | ND | ND | NA |
| OM_D1 | 0 | ND | ND | ND | NA |
|  |  |  |  | **Mean** | **0.37%** |

**Table 4:** Between-Run Reproducibility. Between-run variability expressed as coefficient of variation of mean uantification cycle values of triplicate measurements of all control samples per day. Sample ID includes Pool#)_(<u>H</u>igh/<u>M</u>ed/<u>L</u>ow/<u>N</u>egative/<u>OM</u> buffer only)

| Sample ID | Spike-In Level (fg/μl) | Day 1 (C<sub>q</sub>) | Day 2 (C<sub>q</sub>) | Day 3 (C<sub>q</sub>) | Day 4 (C<sub>q</sub>) | Day 5 (C<sub>q</sub>) | Day 6 (C<sub>q</sub>) | %CV between Runs | %CV between Operators | %CV between Instruments |
| --- | --- | --- | --- | --- | --- | --- | --- | --- | --- | --- |
| P1_H | 1,600,000 | 25.75 | 24.00 | 24.29 | 23.88 | 24.65 | 24.02 | 2.87% | 1.44% | 1.40% |
| P2_H | 1,600,000 | 25.24 | 23.32 | 23.84 | 23.87 | 24.10 | 24.26 | 2.66% | 0.17% | 1.76% |
| P3_H | 1,600,000 | 25.93 | 23.77 | 24.19 | 24.15 | 24.57 | 24.00 | 3.18% | 1.13% | 2.15% |
| P1_M | 160,000 | 28.15 | 27.60 | 28.00 | 27.34 | 28.07 | 27.47 | 1.24% | 0.74% | 0.20% |
| P2_M | 160,000 | 28.01 | 27.65 | 27.32 | 27.11 | 28.09 | 27.16 | 1.55% | 0.53% | 1.14% |
| P3_M | 160,000 | 28.26 | 27.58 | 27.84 | 27.58 | 28.04 | 27.86 | 0.95% | 0.17% | 0.20% |
| P1_L | 5,000 | 34.02 | 33.40 | 33.26 | 33.17 | 33.25 | 33.13 | 0.99% | 0.80% | 0.61% |
| P2_L | 5,000 | 33.56 | 33.23 | 33.32 | 32.99 | 32.64 | 32.62 | 1.15% | 1.33% | 0.57% |
| P3_L | 5,000 | 34.08 | 33.02 | 33.98 | 33.09 | 33.79 | 33.20 | 1.42% | 0.70% | 0.31% |
| P1_N | 0 | ND | ND | ND | ND | ND | ND | NA | NA | NA |
| P2_N | 0 | ND | ND | ND | ND | ND | ND | NA | NA | NA |
| P3_N | 0 | ND | ND | ND | ND | ND | ND | NA | NA | NA |
| OM | 0 | ND | ND | ND | ND | ND | ND | NA | NA | NA |
| **Mean** |  |  |  |  |  |  |  | **1.78%** | **0.78%** | **0.93%** |

Using data generated for between-run reproducibility assessments, we next determined the recovery of the assay by dividing the measured value by the nominal value (the actual amount of DNA that was added to the sample prior to extraction) of the sample (Table 5). The total mean recovery rate of the assay is 41.26% with a standard deviation of 5.74% between measurement days, which met the target acceptance criteria of ≥25%.

**Table 5:** Recovery. Recovery calculated as the percentage of measured to nominal values. Sample ID includes (Pool#)_(<u>H</u>igh/<u>M</u>ed/<u>L</u>ow/<u>N</u>egative/<u>OM</u> buffer only)

| Sample ID | Spike-In Level (fg/μl) | Day 1 | Day 2 | Day 3 | Day 4 | Day 5 | Day 6 | Mean | SD |
| --- | --- | --- | --- | --- | --- | --- | --- | --- | --- |
| P1_H | 1,600,000 | 19.09% | 45.38% | 48.32% | 46.12% | 38.78% | 48.76% | **41.08%** |  |
| P2_H | 1,600,000 | 26.58% | 71.09% | 66.09% | 46.27% | 56.25% | 41.71% | **51.33%** |  |
| P3_H | 1,600,000 | 16.89% | 53.02% | 51.85% | 38.48% | 40.83% | 49.35% | **41.74%** |  |
| P1_M | 160,000 | 39.22% | 42.45% | 39.07% | 46.56% | 39.36% | 48.91% | **42.59%** |  |
| P2_M | 160,000 | 42.89% | 41.27% | 61.54% | 54.20% | 38.78% | 60.07% | **49.79%** |  |
| P3_M | 160,000 | 36.57% | 42.89% | 43.18% | 39.95% | 40.39% | 37.75% | **40.12%** |  |
| P1_L | 5,000 | 26.65% | 29.47% | 34.78% | 31.68% | 39.72% | 35.86% | **33.03%** |  |
| P2_L | 5,000 | 35.86% | 34.97% | 33.23% | 35.67% | 59.22% | 50.76% | **41.62%** |  |
| P3_L | 5,000 | 25.52% | 38.02% | 21.24% | 33.28% | 27.92% | 34.50% | **30.08%** |  |
| P1_N | 0 | ND | ND | ND | ND | ND | ND | **NA** |  |
| P2_N | 0 | ND | ND | ND | ND | ND | ND | **NA** |  |
| P3_N | 0 | ND | ND | ND | ND | ND | ND | **NA** |  |
| OM | 0 | ND | ND | ND | ND | ND | ND | **NA** |  |
| **Mean** |  | **29.92%** | **44.28%** | **44.37%** | **41.36%** | **42.36%** | **45.30%** | **41.26%** | **5.74%** |

To assess potential matrix effects on PCR performance, various spiked stool specimens were compared to spiked OM-200 buffer samples. Matrix interference is reported as the log_10_ deviation between the measurement value of individual non-negative control samples to the mean value obtained from the spiked OM-200 buffer samples. The total mean log_10_ deviation between individual control samples and mean spiked OM-200 buffer was 0.39 log_10_, meeting the target acceptance criteria (Table 6).

**Table 6:**
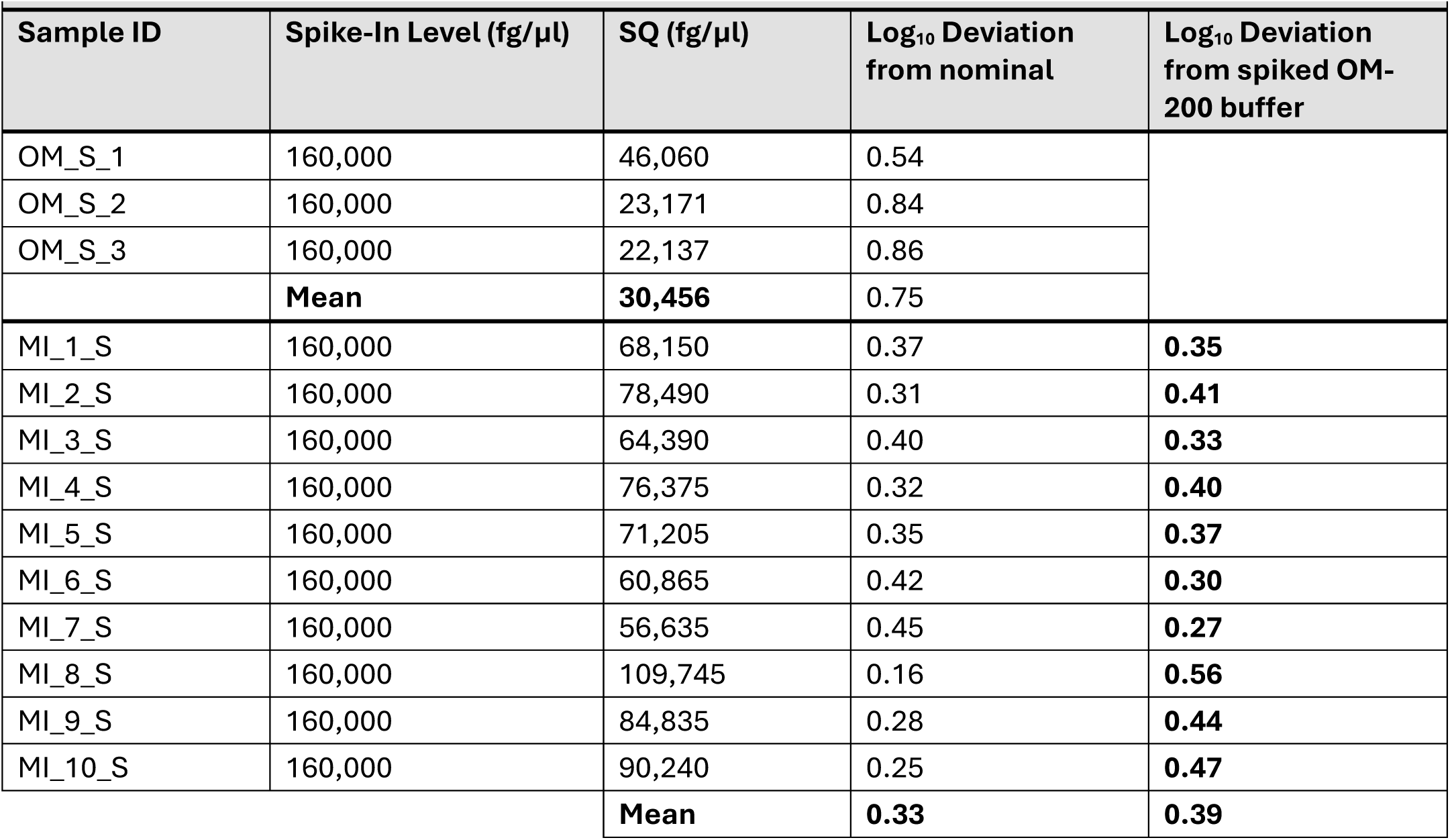
Matrix Interference. Matrix interference expressed as log_10_ deviation between control sample values and mean value of spiked OM-200 buffer samples. OM= OM Buffer, S=Spiked, MI=Matrix terference. #1-10 are individual stool samples that were spiked.

| Sample ID | Spike-In Level (fg/μl) | SQ (fg/μl) | Log<sub>10</sub> Deviation from nominal | Log<sub>10</sub> Deviation from spiked OM-200 buffer |
| --- | --- | --- | --- | --- |
| OM_S_1 | 160,000 | 46,060 | 0.54 |  |
| OM_S_2 | 160,000 | 23,171 | 0.84 |  |
| OM_S_3 | 160,000 | 22,137 | 0.86 |  |
|  | **Mean** | **30,456** | 0.75 |  |
| MI_1_S | 160,000 | 68,150 | 0.37 | **0.35** |
| MI_2_S | 160,000 | 78,490 | 0.31 | **0.41** |
| MI_3_S | 160,000 | 64,390 | 0.40 | **0.33** |
| MI_4_S | 160,000 | 76,375 | 0.32 | **0.40** |
| MI_5_S | 160,000 | 71,205 | 0.35 | **0.37** |
| MI_6_S | 160,000 | 60,865 | 0.42 | **0.30** |
| MI_7_S | 160,000 | 56,635 | 0.45 | **0.27** |
| MI_8_S | 160,000 | 109,745 | 0.16 | **0.56** |
| MI_9_S | 160,000 | 84,835 | 0.28 | **0.44** |
| MI_10_S | 160,000 | 90,240 | 0.25 | **0.47** |
|  |  | **Mean** | **0.33** | **0.39** |

## Discussion

We aimed to establish a reliable, robust, and accurate qPCR assay for detecting *B. infantis* Bi-26 DNA in stool samples, intended for use in clinical research where precise quantification may inform biological insights and potential therapeutic decisions. To ensure the assay’s reliability and suitability for this context, we conducted a comprehensive validation, evaluating key performance parameters including linearity, sensitivity, accuracy, reproducibility, recovery, and matrix interference. This assay demonstrated good performance upon acceptance criteria pre-defined in industry standards, thus demonstrating its suitability for future clinical research applications.

The probiotic market includes a wide range of *B. infantis* strains, both commercially available and under development for therapeutic applications. Different *B. infantis* strains may be evaluated for impact on various health outcomes (23), necessitating an assay that can reliably quantify *B. infantis* regardless of strain variability. Although this validation was conducted using DNA extracted from the *B. infantis* strain Bi-26, the assay targets a conserved genomic region with high sequence identity across many known *B. infantis* strains. The reference standard can be customized to include DNA from the specific strain of interest and be used in a fit-for-purpose validation plan evaluating key parameters necessary for the use-case in future clinical studies. In preliminary evaluations, we observed comparable performance meeting target acceptance criteria when using reference standards prepared from alternative *B. infantis* strains (data not shown). These findings support the assay’s potential utility across a variety of clinical research contexts aiming to evaluate *B. infantis* colonization and its associated health benefits. Furthermore, employing a standardized approach to development and validation of qPCR assays intended to measure *B. infantis* levels in stool across clinical research studies facilitates cross-study comparisons and supports meta-analyses. Such harmonized data can be leveraged to identify threshold levels of total *B. infantis* colonization associated with specific health outcomes. Defining these target colonization levels would be highly valuable for informing the design of future clinical trials, including optimizing dosing strategies and establishing benchmarks for therapeutic efficacy.

This assay has several important limitations that should be considered when interpreting results. First, it was validated using stool samples collected in specialized tubes containing DNA stabilization buffer (OMNIgene® GUT), and it remains unclear whether performance would be equivalent with DNA extracted from flash-frozen stool or samples collected using alternative methods (30, 31). While high-quality DNA is expected to yield comparable results, this was not been formally evaluated in this study. Second, the assay detects *B. infantis* DNA without distinguishing between viable and non-viable bacteria. As such, it may not accurately reflect active colonization by live probiotic *B. infantis* products. This limitation is common in microbiome intervention studies, where assessing bacterial viability through culture-based methods is generally not feasible in probiotic clinical research settings. Therefore, results should be interpreted as representing total *B. infantis* DNA, inclusive of both live and dead cells.

In summary, this study reports a validated qPCR assay for the quantification of *B. infantis* Bi-26 DNA in stool, demonstrating strong performance across key analytical parameters. The assay’s principal suitability for detecting *B. infantis* Bi-26 and potentially additional strains, alongside its applicability with stabilized stool collection methods, make it a valuable tool for clinical research aimed at understanding the role of *B. infantis* in infant health.

## Abbreviations

B Infantis: Bifidobacterium longum subspecies *infantis*
HMOs: Human milk oligosaccharides
WAZ: Weight-for-age Z score
WLZ: Weight-for-length Z score
qPCR: Quantitative real-time PCR
IFF: International Flavors and Fragrances
CV: Coefficient of variation
LLOQ: Lower limit of quantification
ULOQ: Upper limit of quantification
LOD_95_: Limit of Detection (95%)

## Ethics Approval

N/A

## Consent for Publication

N/A

## Data Availability

The datasets used and/or analyzed during the current study are available from the corresponding author on reasonable request.

## Authors’ Contributions

JS, PR, MG, MP, and UZ were involved in the conception or design of validation strategy. MG, MP, AK, and NY contributed to data collection or data generation. MG, JS, MP, PR, NT, AK, NY, HW, UZ, and NF contributed to data analysis or data interpretation. MG, MP, and UZ accessed and verified the data. JS was responsible for the decision to submit the manuscript. All authors reviewed the draft critically, approved the final version to be submitted, and took accountabilities for all aspects of the published work.

## Acknowledgements

The authors thank Loren Blake for their support in assay validation and collaboration coordination as well as Chiara Janssen and Marvin Zumbrunn for technical assistance. Editorial assistance was provided by Madeeha Aqil, PhD, MWC®, CMPP^TM^ and was funded by the Gates Medical Research Institute.

