## Supplement for "A Validated Method for Detection of *Bifidobacterium infantis* Bi-26^TM^ DNA in Stool by Quantitative Real-Time PCR"

**Supplementary Tables**

| Supplementary Table 1: Equipment | |
| --- | --- |
| **Instrument** | **Manufacturer** |
| C1000 Thermal Cycler / CFX96 Optical Reaction Module | Bio-Rad |
| NanoDrop One spectrophotometer | Thermo Scientific |
| Qubit 4 Fluorometer | Invitrogen / Thermo Fisher Scientific |
| Agilent 2100 Bioanalyzer | Agilent Technologies |
| KingFisher Flex System | Thermo Scientific |
| Heated Waterbath – Model 1003 | GFL |
| Centrifuge Rotina 420R | Hettich |
| Centrifuge Micro Star 17 | VWR |
| Centrifuge Universal 320R, Type 1406 | Hettich |
| Precellys Evolution Touch | Bertin |
| 4s 3^TM^ Variable Temperature Thermal Sealer | 4titude |
| Vortexer RS-VA10 | Phoenix Instrument |
| Vortexer MS3 | IKA |

| Supplementary Table 2: Materials | |
| --- | --- |
| Material | Manufacturer |
| TE buffer (10 mM Tris, 0.1 mM EDTA, pH 8.0) | AppliChem (A8569.1000) |
| ZymoBIOMICS^TM^ 96 MagBead DNA Kit | Zymo Research (D4308) |
| Qubit dsDNA HS Assay Kit | Thermo Fisher Scientific (Q32854) |
| Agilent DNA 12000 Kit | Agilent (5067-1508) |
| TaqMan Fast Advanced Master Mix | Thermo Fisher Scientific (4444963) |
| Forward Primer Bi26_F* | Metabion |
| Reverse Primer Bi26_R* | Metabion |
| Probe Bi26_P* | Integrated DNA Technologies, Inc. |
| Nuclease free water | Qiagen (129115) |
| OMNIgene^®^•GUT – OM-200 – Microbiome Stool Self Collection Kit | DNA Genotek Inc. (OM-200) |
| DNA LoBind Tubes 1.5 ml, PCR clean | Eppendorf (0030.108.051) |
| KingFisher 96-deep-well plate (2.0 ml) | LVL (225.DW.2.0.MS.PP) |
| KingFisher 96 (200 μl) microplate | LVL (225.DW.0.2.MS.PP) |
| KingFisher Tip comb for deep-well magnets | LVL (225.DW.TC.MS.PP) |
| Hard Shell PCR Plate | Bio-Rad (HSP9655) |
| Permanent clear heat seal | Bio-Rad (1814035) |

| **Supplementary Table 3: Validation Plan Acceptance Criteria** | |
| --- | --- |
| **Assay Parameter** | **Acceptance Criteria** |
| Dilutional Linearity | R² ≥ 0,98  Efficiency: 90 – 110%  Slope: from -3,1 to -3,6 |
| Sensitivity | NA |
| Accuracy | Quantified value within 1 log_10_ of target value for ≥75% of samples |
| Reproducibility | Within Run:  C_q_ %CV <15% for ≥75% of samples  Between Run:  C_q_ %CV <25% for ≥75% of samples |
| Recovery | Mean Recovery ≥25% |
| Matrix Interference | At least 80% of samples within 1 log_10_ of mean spiked buffer value |
